# Structural insights into specific recognition of PTENα/β-NTE by WDR5 in cancer progression

**DOI:** 10.1101/2023.12.09.570908

**Authors:** Xiaolei Huang, Cheng Zhang, Xinci Shang, Yichang Chen, Qin Xiao, Zhengguo Wei, Guanghui Wang, Xuechu Zhen, Guoqiang Xu, Jinrong Min, Shaoming Shen, Yanli Liu

## Abstract

PTENα/β, two variants of PTEN, play a key role in promoting tumor growth by interacting with WDR5 through their N-terminal extensions (NTEs). This interaction facilitates the recruitment of the SET1/MLL methyltransferase complex, resulting in histone H3K4 trimethylation and upregulation of oncogenes such as *NOTCH3*, which in turn promotes tumor growth. However, the molecular mechanism underlying this interaction has remained elusive. In this study, we determined the crystal structure of PTENα-NTE in complex with WDR5, which reveal that PTENα specifically binds to the WIN site of WDR5 through a novel binding motif specifically found in the NTE domain of PTENα/β. Disruption of this interaction significantly impedes cell proliferation and tumor growth, highlighting the potential of the WIN site inhibitors of WDR5 as therapeutic agents for PTENα/β associated cancers. These findings not only shed light on the important role of the PTENα/β-WDR5 interaction in carcinogenesis, but also present a promising avenue for developing cancer treatments that target this pathway.

---

PTEN (phosphatase and tensin homolog) is a tumor suppressor protein comprised of 403 amino acids, which was initially identified in 1997 by various research groups exploring the cancer susceptibility locus on human chromosome 10q23 (Li and Sun 1997; Li et al. 1997; Steck et al. 1997). Mutations of PTEN have been linked to various cancers, including liver, prostate, and breast cancer (Li and Sun 1997; Li et al. 1997; Steck et al. 1997). Over 80% of Cowden syndrome patients carry PTEN mutations, which are primarily characterized by hamartomas (Yin and Shen 2008). PTEN exhibits both protein phosphatase and lipid phosphatase activities, so it can catalyze the dephosphorylation of p-Tyr, p-Ser, and p-Thr residues in proteins, as well as the phosphoinositide lipid at position D3 of PtdIns(3,4,5)P3, PIP3 (Myers et al. 1997; Maehama and Dixon 1998). By dephosphorylating PIP3, PTEN negatively regulates the PI3K/AKT pathway, which is crucial for cell survival and proliferation. Loss of the PTEN function results in activated PI3K/AKT pathway, promoting cell growth, proliferation, and survival while inhibiting apoptosis (Sulis and Parsons 2003; Sansal and Sellers 2004).

Recent studies have identified multiple non-AUG initiation codons located within its 5’ untranslated region of the PTEN mRNA (Hopkins et al. 2013; Liang et al. 2014; Liang et al. 2017; Zhang et al. 2021). This discovery has led to the identification of various isoforms of the canonical PTEN protein with different lengths of extra amino acids at their N-terminus (Hopkins et al. 2013; Liang et al. 2014; Liang et al. 2017; Zhang et al. 2021). Specifically, three variants known as PTENα (also called PTEN-L) (Hopkins et al. 2013; Liang et al. 2014), PTENβ (Liang et al. 2017), and PTENε (Zhang et al. 2021) have been discovered, which are translated at 519 bp, 438 bp, or 216 bp upstream of the AUG start codon, respectively, and add 173, 146, or 72 amino acids to the N-terminus of the canonical PTEN protein as an N-terminal extension (NTE) domain. Bioinformatic analysis has also revealed that the NTE region contains an intrinsically disordered region (IDR) that is rich in polar residues, potential linear binding motifs, protein-binding sites, and post-translational modification (PTM) sites. These findings suggest that the NTE may serve as a signaling platform in regulating the biological functions and intracellular trafficking of PTEN (Malaney et al. 2013).

The initial study on PTENα revealed that it possesses a secretion signal sequence and a resembled cell-penetrating element, which enable it to be secreted out of cells and taken up by other cells. This ability allows PTENα to inhibit the growth of glioblastoma tumors by interfering with the PI3K signaling pathway (Hopkins et al. 2013). Meanwhile, another study demonstrated that PTENα is localized in mitochondria, where it regulates energy metabolism (Liang et al. 2014). In contrast, PTENβ that was initially found in the nucleus negatively regulates nucleolin phosphorylation levels, pre-RNA synthesis and cell prolilferation (Liang et al. 2017). On the other hand, recent research has shown that both PTENα and PTENβ (without the secretion signal sequence) can be secreted into the extracellular space and cleaved by the proprotein convertase Furin, yielding a C-terminal fragment that significantly suppresses the proliferation of tumor cells (Zhang et al. 2022). Besides being present in the extracellular space, PTENα and PTENβ are also prominently localized in the nucleus and promote liver cancer cell growth by interacting with WDR5 protein through their NTEs (Shen et al. 2020). PTENε, a smaller PTEN variant primarily located in the cell plasma membrane, inhibits the formation of pseudopods and reduces the migratory ability of tumor cells, thereby repressing cancer cell metastasis (Zhang et al. 2021). Overall, these findings emphasize the crucial roles played by PTENα/β/ε in maintaining normal cell survival and inhibiting or promoting tumor cell proliferation and migration, underscoring the complexity of their involvement in tumor progression.

WDR5 is a highly conserved protein with a seven-bladed β-propeller fold. It is an essential subunit of the SET1/MLL methyltransferase complexes, which regulate gene expression by catalyzing methylation at histone H3K4 sites (Schuettengruber et al. 2011; Shilatifard 2012). WDR5 performs its role by presenting H3K4 for methylation, through the recognition of the guanidyl group of H3R2 by utilizing its central channel of the β-propeller structure located at the top face (Schuetz et al. 2006). This region is known as WDR5 interacting site or the WIN site (Patel et al. 2008). Apart from H3R2, WDR5 also interacts with other arginine-containing sequences called the WIN motif from the catalytic component of the SET1/MLL complexes (Patel et al. 2008; Dharmarajan et al. 2012; Zhang et al. 2012) and other proteins (Guarnaccia and Tansey 2018). Due to its unique epigenetic role by means of interacting with various ligands, WDR5 is involved in many biological processes, including reproduction (Jiang et al. 2009; Xu et al. 2012), development (Wysocka et al. 2005; Zhu et al. 2008; Diao et al. 2012), metabolism (Ravnskjaer et al. 2013), immune responses and inflammation (Wang et al. 2010; Gagnon and Corey 2012; Tennant et al. 2015), neural and humoral regulation (Ka et al. 2022); (Mo et al. 2006). Additionally, WDR5 is implicated in the onset, progression, and maintenance of multiple diseases. For instance, overexpression of WDR5 is not only associated with the development and progression of various cancers, such as prostate cancer (Kim et al. 2014; Kim et al. 2016), breast cancer (Dai et al. 2015), leukemia (Ge et al. 2016), and liver cancer (Cui et al. 2018), but also linked to unfavorable clinical outcomes. Consequently, it appears to be a promising approach for treatment of these diseases by targeting WDR5.

The previous study has shown that PTENα/β promotes tumor progression in liver cancer cells by interacting with WDR5 to recruit the SET1/MLL complexes to the promoters of the PTENα/β-target genes, such as *NOTCH3*, a broadly studied oncogene (Ali et al. 2016; Jeong et al. 2017; Shen et al. 2020). Therefore, understanding the molecular mechanism of the interactions between PTENα/β and WDR5 and deciphering its underlying molecular functions hold substantial importance for exploring potential therapeutic applications. In this study, we aimed to investigate the structural details of the interactions between PTENα/β-NTE and WDR5 using biophysical binding assays and X-ray crystal structure analysis. We discovered that PTENα/β-NTE utilizes an N-terminal WIN motif to interact with the arginine-binding pocket of WDR5. Interestingly, we identified a unique-RR-motif in PTENα/β-NTE for binding to the WDR5 WIN site, which differs from the conserved-A/CR-motif found in other WIN motif sequences. To validate the functional significance of this interaction, cell and xenograft mouse experiments were conducted, which demonstrated that disrupting the PTENα-WDR5 interaction reduces the expression of downstream oncogenes, inhibiting cell proliferation and tumor growth. Our findings provide valuable insights into the crucial role of PTENα/β-NTE in cancer biology and offer potential therapeutic targets for cancer treatment.

## Results

### The N-terminal extension (NTE) of PTENα/β binds to WDR5

A previous study has revealed that PTENα/β-NTE requires two crucial motifs for their tumorigenic activity, a WDR5 binding motif (amino acids 116-148) and an essential nuclear localization signal (NLS) (Fig. 1a) (Shen et al. 2020). To confirm the interaction between PTENα/β-NTE and WDR5, isothermal titration calorimetry (ITC) assays were conducted to determine the dissociation constants (*K_d_*) of WDR5^22-334^ with a synthetic PTENα-NTE^116-148^ peptide and the recombinant PTENα-NTE^1-173^ protein. The ITC data indicated that both the PTENα-NTE^116-148^ peptide and the recombinant PTENα-NTE^1-173^ protein could bind to WDR5^22-334^, with the former exhibiting a higher binding affinity than the latter (*K_d_*: 3.8 μM *vs.* 17 μM, Fig. 1b). This suggested that the PTENα-NTE^116-148^ fragment is sufficient for their interaction *in vitro*. Our ITC data provide further evidence supporting the fact that PTENα/β-NTE directly binds to WDR5 *in vitro,* since the PTENα-NTE^116-148^ fragment is a common region for both NTEs of the two PTEN variant proteins.

**Figure 1.**
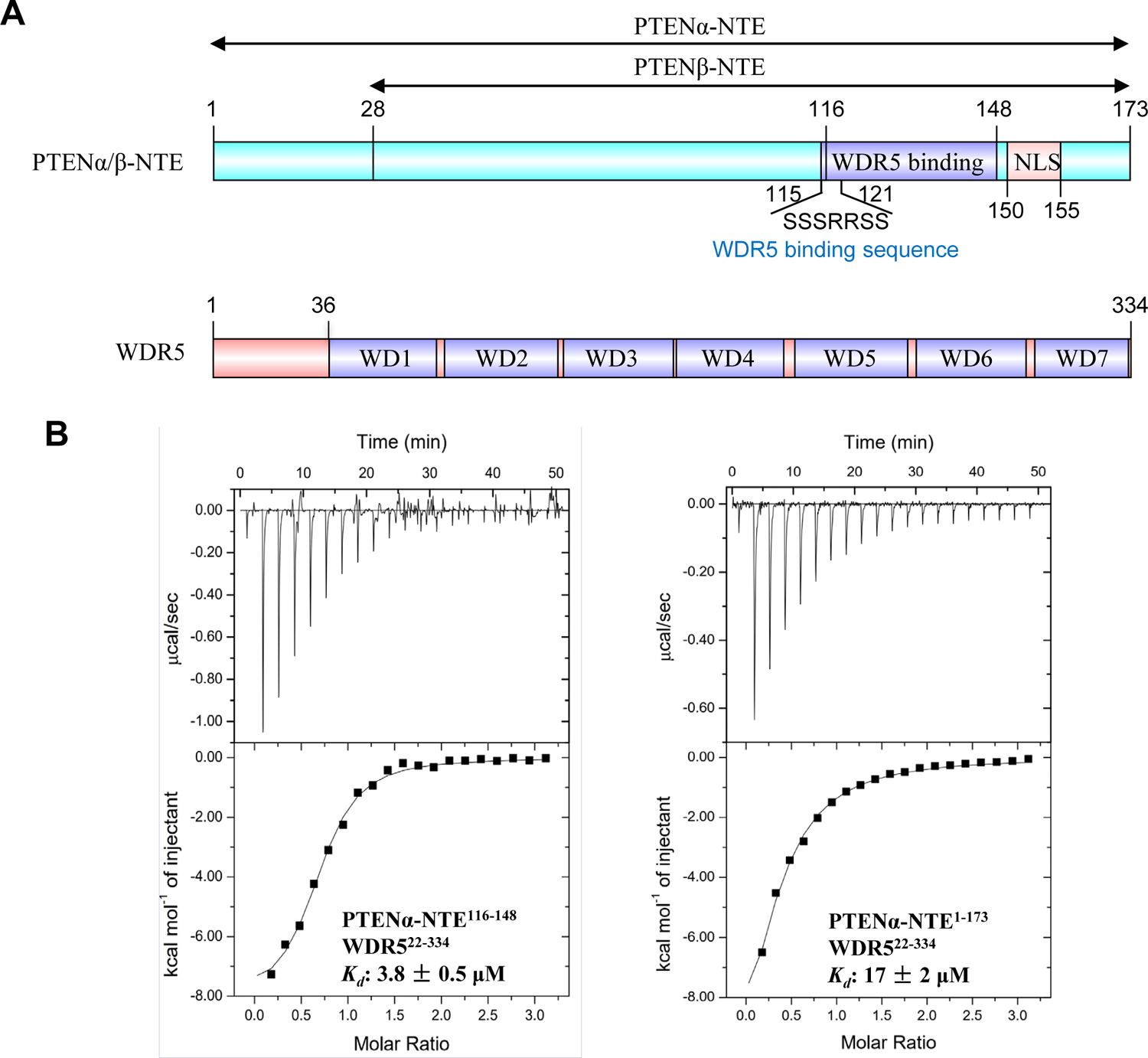
PTENα/β-NTE binds to WDR5. (*A*) Domain structure of human PTENα/β-NTE and WDR5. PTENα/β-NTE consists of a WDR5 binding motif, a nuclear localization signal (NLS). WDR5 consists of seven WD40 repeats. (*B*) ITC binding curves for the titration of PTENα-NTE^116-148^ or PTENα-NTE^1-173^ to WDR5^22-334^ by iTC-200 microcalorimeter (MicroCal, Inc.). *K_d_*: dissociation constants (μM); ITC data shown are representative of two independent experiments and all *K_d_* values were calculated from single measurement and errors were estimated by fitting curve.

### PTENα/β-NTE binds to the WIN site of WDR5 by using an -RR-motif

To better understand how PTENα/β and WDR5 interact at the molecular level, we attempted to crystallize WDR5 in complex with PTENα-NTE or the synthetic PTENα-NTE^116-148^ peptide and successfully solved the crystal structures of WDR5 bound and unbound by PTENα-NTE^1-173^ (Fig. 2, Supplementary Fig. 1 and Table 1). Initially, we attempted to co-crystallize the synthetic PTENα-NTE^116-148^ peptide with full-length WDR5 (amino acids 1-334), but only obtained the crystal structure of WDR5 in the PTENα-NTE peptide-free form at 1.8 Å resolution. In this apo WDR5 structure, the N-terminal residues (^11^EAARAQPT^18^) of WDR5 occupy the WIN site, with WDR5-R14 inserting into the negatively charged channel of the WIN site, as previous studies have shown (Schuetz et al. 2006; Tan et al. 2021) (Supplementary Fig. 1). Therefore, we tried using a WDR5^22-334^ truncation for co-crystallization screening with PTENα-NTE^116-148^ or PTENα-NTE^1-173^ and eventually obtained the co-crystals of WDR5^22-334^ with PTENα-NTE^1-173^.

**Figure 2.**
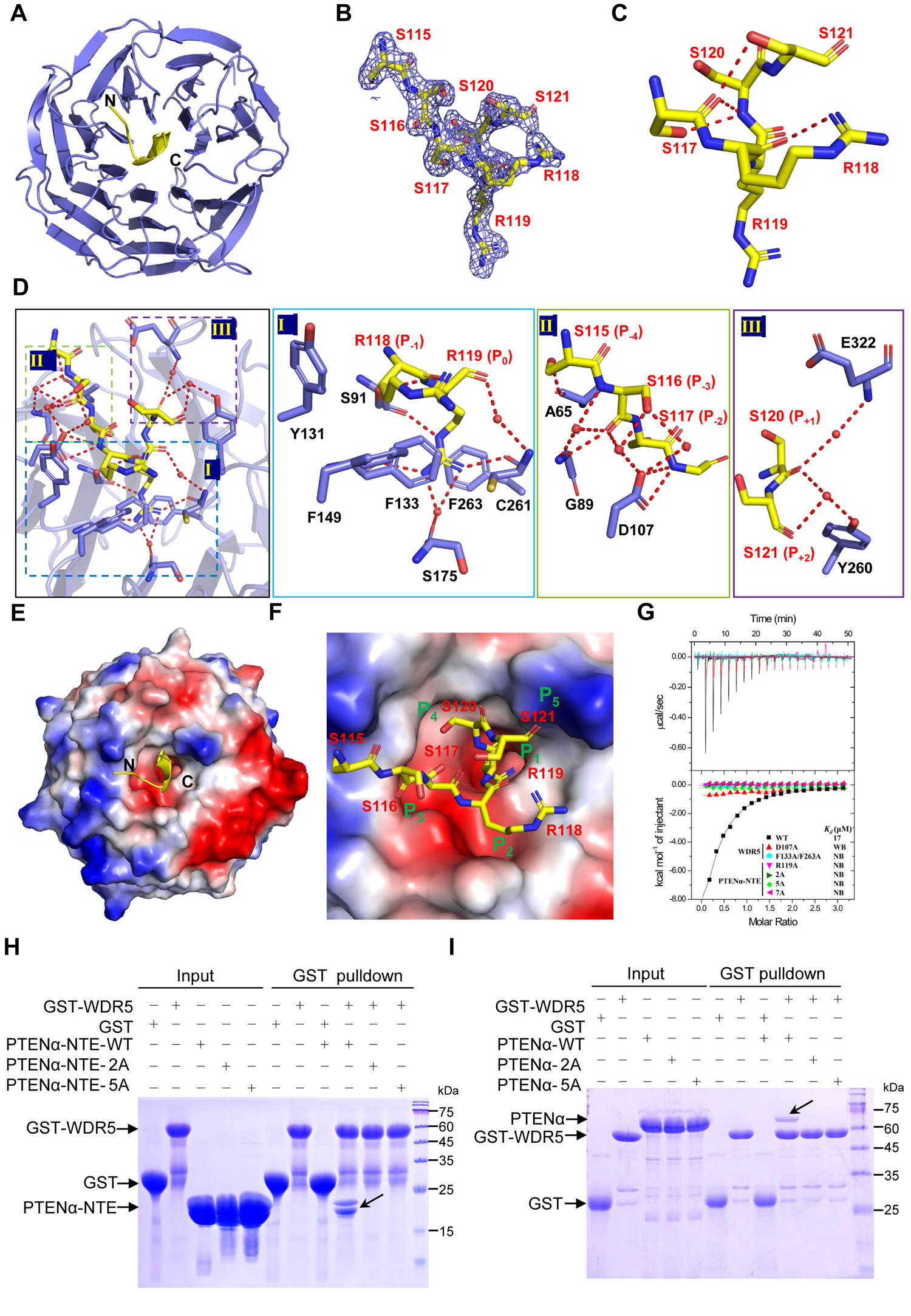
Crystal structure of WDR5^22-334^ in complex with PTENα-NTE^1-173^. (*A*) Overall structure of WDR5 in complex with PTENα-NTE. The structure was shown in cartoon with WDR5 colored in slate and PTENα-NTE colored in yellow. (*B*) Fo-Fc omit map of PTENα-NTE contoured at 1σ level. (*C*) Intramolecular hydrogen bonds stabilize the WDR5-PTENα-NTE interaction. (*D*) Detailed interaction between PTENα-NTE and WDR5. Amino acid residues of WDR5 involved in the PTENα-NTE interaction were shown as sticks and the detailed interactions were shown by enlarged views. Key hydrogen bonds were depicted as red dash lines and key water molecules were indicated as red sphere. (*E, F*) Electrostatic potential surface view of WDR5 in complex with PTENα-NTE. The PTENα-NTE was shown as cartoon (*E*) and sticks (*F*), respectively. Five WIN motif binding pockets (P_1_-P_5_) in WDR5 were labeled with green (*F*). (*G-I*) Mutation of the interacting residues affected the interaction between PTENα-NTE^1-173^ or full-length PTENα^1-576^ and WDR5^22-334^. ITC binding curves for the titration of different mutants of PTENα-NTE to wild-type WDR5 or wild-type PTENα-NTE to mutant WDR5 by iTC-200 microcalorimeter (MicroCal, Inc.) (*G*). *In vitro* GST pulldown of WDR5 with wild-type or mutant of PTENα-NTE (*H*) or full-length PTENα (*I*). Bacterially expressed proteins PTENα-NTE (*H*) or full length PTENα (*I*) and their mutants, as indicated, were incubated with GST or GST-tagged WDR5, followed by GST pulldown and CBB staining, with the specific binding indicated by an arrow. *K_d_*: dissociation constants (μM); NB: no detectable binding; WB: weak binding; 2A: PTENα-NTE_R118A/R119A; 5A: PTENα-NTE_115-119-5A; 7A: PTENα-NTE_115-119-5A_R135A/R144A.

**Table 1.** Data collection and refinement statistics.

|  | WDR5-PTEN | WDR5 |
| --- | --- | --- |
| **PDB code** | 8X3S | 8X3R |
| **Data collection** |  |  |
| Space group | P 2<sub>1</sub> 2<sub>1</sub> 2<sub>1</sub> | P 2<sub>1</sub> 2<sub>1</sub> 2<sub>1</sub> |
| Cell dimensions |  |  |
| *a*, *b*, *c* (Å) | 46.5, 63.8, 92.8 | 48.6, 52.7, 123.1 |
| $\alpha$, $\beta$, $\gamma$ (°) | 90, 90, 90 | 90, 90, 90 |
| Resolution (Å) | 29.19-1.87 (1.91-1.87) | 24.21-1.76 (1.79-1.76) |
| Measured reflections | 40296 (1831) | 55450 (2918) |
| Unique reflections | 22704 (1171) | 30780 (1719) |
| *R*<sub>merge</sub> | 0.071 (0.533) | 0.045 (0.428) |
| *I*/σ*I* | 10.5 (1.8) | 10.9 (1.7) |
| CC<sub>1/2</sub> | 0.994 (0.627) | 0.998 (0.674) |
| Completeness (%) | 97.0 (80.3) | 96.9 (96.6) |
| Redundancy | 1.8 (1.6) | 1.8 (1.7) |
| **Refinement** |  |  |
| Resolution (Å) | 29.19-1.87 (1.94-1.87) | 24.21-1.76 (1.82-1.76) |
| *R*<sub>work</sub>/*R*<sub>free</sub> (%) | 22.4/25.7 | 19.7/21.2 |
| No. of atoms/average *B*-factors (Å<sup>2</sup>) | 2558/33.2 | 2604/19.9 |
| Protein | 2377/32.6 | 2350/19.6 |
| Ligand | 52/36.1 | 65/23.2 |
| Water | 129/37.0 | 189/20.0 |
| R.m.s. deviations |  |  |
| Bond lengths (Å) | 0.01 | 0.01 |
| Bond angles (°) | 1.3 | 1.2 |

Values in parentheses are for the highest resolution shell.

The complex structure revealed that WDR5 exhibits the typical seven-bladed β-propeller structure, while PTENα-NTE adopts a 3_10_-helical conformation fitting into the central WIN site pocket of WDR5 (Fig. 2a). Based on the electron density, the residues 115 to 121 of PTENα-NTE could be distinctly traced and form numerous intramolecular hydrogen bonds to compact its conformation (Fig. 2b-c). The interaction between PTENα-NTE and WDR5 involves a network of hydrogen bonds, van der Waals contacts, hydrophobic packing, and cation-π interactions, which are observed in other WDR5 complexes involving the accommodation of the WIN motif ligands within the WIN site (Fig. 2d-f) (Schuetz et al. 2006; Dharmarajan et al. 2012; Zhang et al. 2012). Specifically, the key R119 (P_0_) residue of PTENα-NTE extends into the pocket 1 of WDR5, interacting with F133 and F263 side chains through cation-π interaction, as well as making several direct or water-mediated hydrogen bonds with the main chains of S91, F133, S175, and C261 to further stabilize the interaction (Fig. 2d-Ⅰ and 2f). While the PTENα-R118 (P_-1_) residue is embraced by pocket 2 via hydrophobic packing with Y131 and F149 residues of WDR5 (Fig. 2d-Ⅰ and 2f). The S115-S117 (P_−4 to −2_) residues of PTENα-NTE bind to pocket 3 and form several main-chain and side-chain hydrogen bonds with A65, G89, and D107 residues of WDR5 (Fig. 2d-Ⅱ and 2f). The C-terminus of the PTENα-NTE^115-121^ fragment, comprising S120 (P_+1_) and S121 (P_+2_) residues, is held by pockets 4 and 5 through direct or water-mediated hydrogen bonds with E322 and Y260 residues of WDR5 (Fig. 2d-Ⅲ and 2f). The PTENβ-NTE is expected to bind to WDR5 in the same way as PTENα-NTE since PTENα-NTE^115-121^ fragment is a common region for both of them. Overall, our complex structural studies demonstrated that PTENα/β-NTE binds to the WIN site of WDR5 snugly by forming numerous intermolecular interactions.

### PTENα/β-NTE interacts with WDR5 exclusively through the WIN site

To verify our findings from the complex crystal structure, we introduced specific point mutations into WDR5^22-334^ for binding studies. The introduction of the double point mutations WDR5_F133A/263A (referred to as WDR5-2A) into WDR5^22-334^ resulted in the complete loss of the interaction between WDR5 and PTENα-NTE (Fig. 2g, Supplementary Fig. 2a), which is consistent with the crucial role played by these two residues in WIN site binding, as indicated by previous studies (Couture et al. 2006; Ruthenburg et al. 2006; Song and Kingston 2008). Additionally, when we introduced the D107A mutation into WDR5^22-334^, another critical residue for WIN site binding, the capacity of WDR5^22-334^ to bind to PTENα-NTE^1-173^ was significantly reduced (Fig. 2g, Supplementary Fig. 2a), which also aligns with previous studies (Couture et al. 2006; Ruthenburg et al. 2006; Song and Kingston 2008). Collectively, all these mutants binding assays confirmed that WDR5 recognizes PTENα/β-NTE specifically through the WIN binding site.

To further confirm whether PTENα-NTE^1-173^ interacts with WDR5 only through the WIN binding site, a competitive binding assay was conducted using unmodified histone H3 peptide with residues 1-15 (H3^1-15^), a well-known ligand of the WDR5 WIN site (Ruthenburg et al. 2006; Schuetz et al. 2006). ITC experiments were performed with PTENα-NTE^116-148^ and PTENα-NTE^1-173^ against a mixture of WDR5^22-334^ and H3^1-15^ with a 1:1 molar ratio. The results revealed that PTENα-NTE^116-148^ weakly bound to the WDR5^22-334^ and H3^1-15^ mixture, with a *K_d_* of 32 ± 3 μM. However, no significant binding tendency was observed for PTENα-NTE^1-173^, which may be due to PTENα-NTE^116-148^ having a higher binding affinity for WDR5^22-334^ than H3^1-15^, while PTENα-NTE^1-173^ and H3^1-15^ having a comparable binding affinity for WDR5^22-334^ (*K_d_* values: 3.8 ±0.5, 17 ±2, and 25 ±3 μM for PTENα-NTE^116-148^, PTENα-NTE^1-173^, and H3^1-15^, respectively, Supplementary Fig. 2b-c). Reverse competitive titration experiments using H3^1-15^ against a mixture of WDR5^22-334^ with PTENα-NTE^1-173^ or PTENα-NTE^116-148^ were also conducted, and no notable binding was observed (Supplementary Fig. 2c). Overall, all these binding assays support the conclusion that PTENα/β-NTE interacts with WDR5 exclusively through the WIN site.

### PTENα/β-NTE interacts with the WDR5 WIN site solely through the SSSRRSS WIN motif

To investigate the significance of the ^115^SSSRRSS^121^ WIN motif within PTENα-NTE^1-173^ in binding to wild-type WDR5^22-334^, a series of ITC assays were conducted using various PTENα-NTE^1-173^ mutants. Initially, experiments were carried out using 6×His-TEV (tobacco etch virus)-PTENα-NTE^1-173^ fusion proteins for ease of purification. Previous reports have highlighted the arginine residue as a major structural determinant for the WDR5 binding (Schuetz et al. 2006; Dharmarajan et al. 2012). Therefore, to avoid any possible compensatory effect between R118 and R119 (Supplementary Fig. 3a), a double point mutant (PTENα-NTE^1-173^-2A: PTENα-NTE^1-173^_R118A/R119A) was tested, which displayed weak binding to WDR5^22-334^ (Supplementary Fig. 3b). Furthermore, the PTENα-NTE^1-173^_115-119-5A mutant (PTENα-NTE^1-173^-5A), and the PTENα-NTE^1-173^-5A_R135A/R144A (PTENα-NTE^1-173^-7A) mutant with mutation of all the arginine residues within PTENα-NTE^116-148^ were generated, which still displayed weaker interactions with WDR5 (Supplementary Fig. 3a-b).

Subsequent literature research identified that the arginine residue within the 6×His-TEV-tag can mimic the WIN motif and bind to the WDR5 WIN site (Schuetz et al. 2006), explaining why the tagged PTENα-NTE^1-173^-7A mutant still retained some binding to WDR5. Therefore, all mutant ITC assays were repeated with the His-TEV-tag removed proteins, and the results revealed that none of the mutants bound to WDR5^22-334^ (Fig. 2g, Supplementary Fig. 3c). Moreover, even a single PTENα-NTE^1-173^-R119A mutant was sufficient to abolish its binding to WDR5^22-334^ (Fig. 2g, Supplementary Fig. 3c), indicating that the R119 of PTENα is critical for the interaction between PTENα/β-NTE and WDR5. Finally, the GST pulldown assay demonstrated that WDR5^22-334^ directly and specifically bound to both wild-type PTENα-NTE^1-173^ and full length PTENα^1-576^ proteins *in vitro*, but not the mutants containing the R119A (Fig. 2h-i). These findings strongly support the conclusion that PTENα/β-NTE interacts with WDR5 WIN site solely through the SSSRRSS WIN motif.

### A novel -RR-binding motif was identified in PTENα/β-NTE for binding to WDR5

Comparison with the published WDR5-WIN motif complex structures revealed that all the WIN motifs bind to WDR5 similarly through its conserved arginine residue inserting into the central tunnel of WDR5 and the residue at P_-1_ position displays a preference for a small side-chain residue (Fig. 3a and Supplementary Fig. 4). However, the WIN motif of PTENα/β-NTE, namely the SSSRRSS fragment, bears a large side-chain residue, arginine, at the P_-1_ position, different from the previous reported WIN site ligands with smaller side-chain residues, such as alanine (Dharmarajan et al. 2012; Zhang et al. 2012; Dias et al. 2014; Qin et al. 2014; Guarnaccia et al. 2021) and cysteine (Tan et al. 2021) (Fig. 3a-b and Supplementary Fig. 5). The side chain of R118 of PTENα-NTE^1-173^ extends into another pocket (P_2_ pocket) and forms hydrophobic packing with Y131 and F149 side chains of WDR5 (Fig. 3c-Ⅰ). To investigate the impact of amino acid identity at position P_-1_ on binding affinity, ITC experiments were performed using various PTENα-NTE^1-173^-R118 mutants (Fig. 3d and Supplementary Fig. 6). As shown in Fig. 3d, all the mutants except R188A weakened the binding due to steric hindrance or lack of enough hydrophobic interaction. Among these mutants, mutating arginine to lysine, which has a similar size and charge to arginine, resulted in weak binding to WDR5^22-334^ too. Further careful structural analysis revealed that an intramolecular hydrogen bond between the side-chain guanidyl group and the main-chain carbonyl group of R118 allows the side chain of R118 to orient closer to the main chain, eliminating spatial hindrance of its insertion into the P_2_ pocket (Fig. 3c-Ⅱ). Consequently, the bent side chain of R118 finds a perfect fit within the P_2_ pocket (Fig. 3c-Ⅲ), while the side chain of lysine could not form this intramolecular hydrogen bond and may cause spatial hindrance. Overall, these structural and binding studies highlight the importance of the amino acid identity at the P_-1_ position to the binding, and the novel finding that an arginine residue at the P_-1_ position exhibits robust binding affinity will broaden the ligand repertoire, providing the chance to search for additional WIN site binding partners.

**Figure 3.**
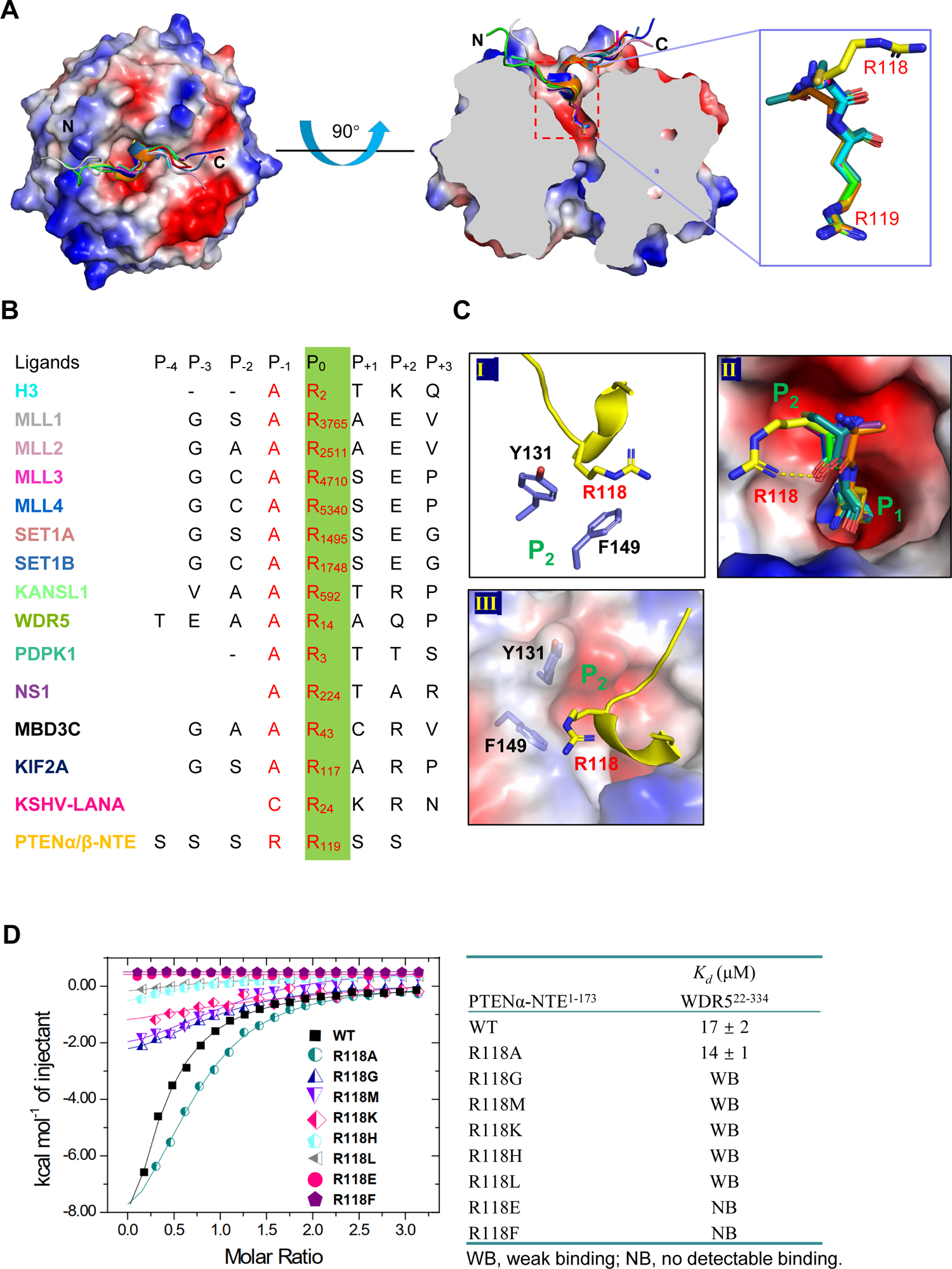
Structural comparison with other WIN site ligands. (*A*) Different WIN motif ligand peptides bind to WDR5 using the same arginine-binding pocket. Superposition of WIN motif peptides shown with a schematic representation (*left*), cut-away view of the arginine-binding pocket of WDR5 (*middle*), and zoomed view of the conserved WIN motif residues (P_0_ and P_-1_) (*right*). The WDR5 molecule was presented as electrostatic potential surface and the different WIN motif peptides were presented as cartoon diagram by different colors. (*B*) PTENα/β-NTE has a unique -RR-WIN site binding motif. Sequence alignment of the WIN motif peptides displayed in the complex structures. The key conserved arginine residue was highlighted in green and the conserved WIN motif residues (P_0_ and P_-1_) were represented in red character. (*C*) The unique R118 residue of the PTENα-NTE binds to the P_2_ pocket of WDR5. (*D*) Mutations of the PTENα-NTE-R118 residue affected binding to WDR5. ITC curves (*left*) and binding affinities (*right*) for the titration of wild-type or different R118 mutants of PTENα-NTE^1-173^ to WDR5^22-334^.

### PTENα stimulates gene expression through its interactions with WDR5

To confirm that PTENα interacts with WDR5 through its SSSRRSS motif in cell, 3×FLAG-tagged WDR5 was co-transfected with either wild-type HA-tagged PTENα or its 5A and 7A mutants into HEK293T cells. Co-IP assays using an anti-FLAG antibody disclosed that only PTENα was successfully co-immunoprecipitated with WDR5, while PTENα-5A and PTENα-7A did not show any interaction with WDR5 (Fig. 4a). These results were consistent with our findings from the complex structure, ITC and *in vitro* GST pulldown data, highlighting the critical role of the PTENα-NTE SSSRRSS motif in interacting with WDR5 in a cellular context.

**Figure 4.**
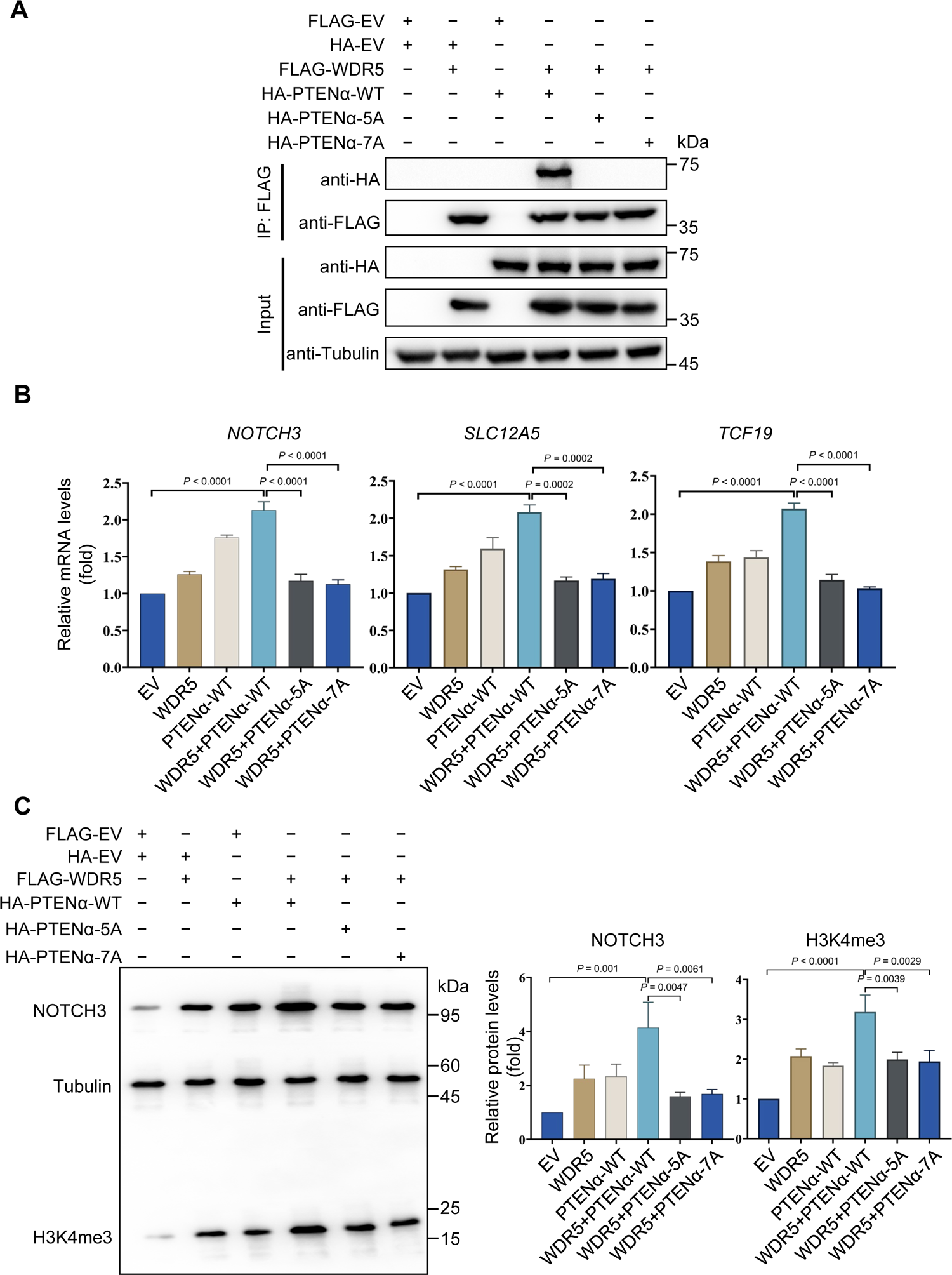
The interaction between PTENα-NTE and WDR5 plays a crucial role in enhancing the transcriptional activity and expression of genes associated with tumorigenicity. HEK293T cells were transfected with 3×FLAG-tagged WDR5 along with wild-type PTENα, PTENα-5A, or PTENα-7A mutants. Subsequently, a series of experiments were conducted: (*A*) Mutation of the key interacting residues of PTENα-NTE affected the interaction in cells. Western blotting was performed on the indicated proteins in co-immunoprecipitates (Co-IPs). (*B*) Disruption of the interaction between PTENα and WDR5 impaired transcriptional activity of target genes, such as *NOTCH3*, *SLC12A5*, and *TCF19* determined by qRT-PCR. (*C*) Disruption of the interaction between PTENα and WDR5 decreased the protein level of NOTCH3 and trimethylated histone H3K4 determined by western blotting. The experiments were repeated three times independently with similar results, and the results of one representative experiment were shown. For *B-C*, data represent means ±s.e.m. Statistical significance was determined by two-tailed unpaired *t*-test. For *A*, *C*, pre-stained protein marker: Abclonal, RM19001.

Previous study has shown that the interaction between PTENα/β and WDR5 plays a crucial role in recruiting the SET1/MLL methyltransferase complex to regulate the downstream H3K4me3 level and expression of oncogenes such as *NOTCH3*, *SLC12A5*, and *TCF19* (Shen et al. 2020). To validate the importance of the SSSRRSS motif of PTENα-NTE in regulating the expression of these oncogenes and the H3K4me3 level, qRT-PCR and western blot experiments were conducted. The results demonstrated that co-expression of WDR5 and wild-type PTENα had a more potent effect compared to the expression of either WDR5 or wild-type PTENα on the mRNA levels of *NOTCH3*, *SLC12A5*, as well as *TCF19*, and the levels of H3K4me3 modification and NOTCH3 (Fig. 4b-c). However, this enhanced effect was disrupted in the mutant of the SSSRRSS motif, which showed comparable effects to that of WDR5 expression alone (Fig. 4b-c). Overall, these results confirm the critical role of the SSSRRSS motif of PTENα-NTE in interacting with WDR5, as well as modulating downstream epigenetic factors and gene expression related to tumorigenicity.

### PTENα promotes tumorigenesis through the interaction between the PTENα-NTE SSSRRSS motif and WDR5

It has been previously shown that PTENα contributes to the tumor development of SMMC-7221 cancer cells (Shen et al. 2020). To investigate the role of the SSSRRSS motif of PTENα-NTE in promoting tumorigenesis of these cells, mutants of PTENα-5A/7A were used to examine whether they could impede PTENα-mediated tumorigenesis promotion in the stable *PTEN*3 (PTEN, PTENα, and PTENβ) knockout cell line (SMMC-7721-*PTEN*3^KO^). Ectopic expression of wild-type PTENα, but not PTENα-5/7A, restored the pro-proliferative effect in these cells, while the ectopic expression of PTEN inhibited cell proliferation, consistent with its function as a tumor suppressor (Fig. 5a-c). Additionally, ectopic expression of PTENα had no impact on AKT activation in *PTEN*3^KO^ SMMC-7721 cells, consistent with previous study (Shen et al. 2020) (Fig. 5a). When these cells were subcutaneously inoculated into nude mice, the ectopic expression of wild-type PTENα accelerated tumorigenesis, while the PTENα-5A/7A groups did not show significant difference from the control group (Fig. 5d-e).

**Figure 5.**
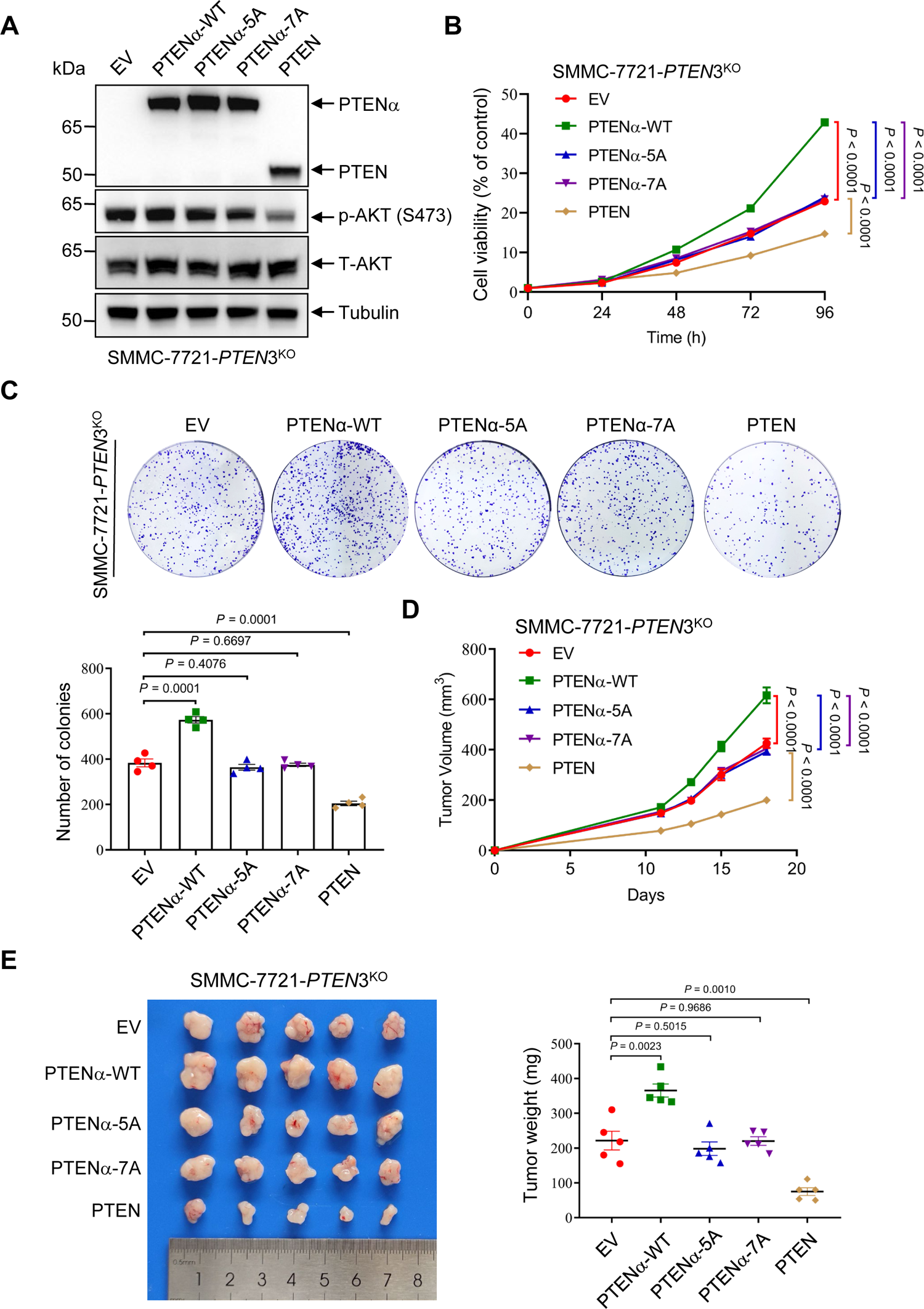
PTENα promotes tumorigenesis through the interaction between WDR5 and PTENα-NTE. (*A*) The efficiency of rescue of *PTEN3*^KO^ SMMC-7721 cells by ectopic expression of wild-type PTEN, PTENα or its mutants. SMMC-7721 *PTEN3*^KO^ cells were transduced by lentiviruses encoding wild-type PTEN, PTENα or its mutants, followed by western blotting of the indicated proteins. Pre-stained protein marker: ThermoFisher, 26616. (*B, C*) Disruption of the interaction between PTENα and WDR5 impaired tumor cell growth and tumorigenic capacity. CCK8 assays (*B*) and colony formation assays (*C*) were used to determine the proliferation and tumorigenic capacity of these cells. (*D, E*) Disruption of the interaction between PTENα and WDR5 destroyed the promotion of tumorigenesis by PTENα. The transfected cells were subcutaneously injected into nude mice (1×10^6^ cells per mouse; n = 5 mice per group). Tumor volumes were measured at different days (*D*). On day 17, tumors were harvested, photographed (*left*), and weighed (*right*) (*E*). The experiments were repeated three times (twice for animal experiments) independently with similar results, and the results of one representative experiment were shown. For *B-E*, data represent means ±s.e.m. Statistical significance was determined by two-way ANOVA (*B, D*) or two-tailed unpaired *t*-test (*C, E*). T-AKT, total AKT.

To further investigate the role of WDR5 in cancer progression, a *WDR5* knockout SMMC-7721 cell line (SMMC-7721-*WDR5*^KO^) was generated and rescued it with empty vector (EV), wild-type WDR5, or WDR5-2A mutant (F133A/F263A) (Supplementary Fig. 7a). The results revealed that knocking out *WDR5* significantly reduced cell proliferation and carcinogenesis, whereas ectopic expression of wild-type WDR5 promoted potent cell proliferation and tumor growth in comparison to all other cell lines (Supplementary Fig. 7b-e). Notably, ectopic expression of the WDR5-2A mutant partially rescued cell proliferation and the tumor-promoting effect owing to the retention of the WBM (WDR5-binding motif) site, which interacts with other proteins such as MYC (Thomas et al. 2015). These findings suggest that the interaction between the SSSRRSS motif of PTENα and WDR5 is indispensable for PTENα-mediated cancer cell proliferation and tumor growth. However, this interaction is not the only requirement for pro-tumor activity of WDR5. Interactions of WDR5 with other proteins such as MYC may also play a role in promoting tumor growth. Overall, these results provide new insights into the mechanisms underlying cancer cell proliferation and tumor growth, highlighting potential targets for developing new therapeutic strategies.

## Discussion

In this study, we identified a novel binding motif, SSSRRSS, in the PTENα/β-NTE sequence that interacts with the WDR5 WIN site. The WDR5 WIN site, one of the two well-known binding sites of WDR5, has been found to interact with various ligands, exerting different physiological functions, many of which are linked to cancer development (Kim et al. 2014; Dai et al. 2015; Ge et al. 2016; Cui et al. 2018). Typically, this site recognizes its ligands by enveloping the conserved arginine residue of the ligands by its central negatively charged channel and accommodating a small side-chain residue preceding the conserved arginine residue (P_-1_ position), such as alanine (Dharmarajan et al. 2012; Zhang et al. 2012; Dias et al. 2014; Qin et al. 2014; Guarnaccia et al. 2021) or cysteine (Tan et al. 2021), via a shallow surface pocket. However, a new -RR-WIN site binding motif was discovered that features a larger side-chain arginine at the P_-1_ position. Structural analysis revealed that a specific intramolecular hydrogen bond of the P_-1_ arginine stabilizes its side chain, significantly reducing steric hindrance to allow for proper interaction with WDR5. These findings provide new insights into the structural determinants governing the interaction between PTENα/β-NTE and WDR5, as well as redefine the sequence characteristics of the WDR5 WIN site ligands.

A previous study has suggested that PTENα and PTENβ contribute to tumor progression by interacting with WDR5 (Shen et al. 2020). Our structural and binding studies demonstrated that a common fragment of PTENα/β, SSSRRSS, specifically binds to the WIN site of WDR5 and the mutations of the key interaction residues disrupt this association significantly. Furthermore, our *in vitro* and *in vivo* studies revealed that the SSSRRSS fragment is indispensable for the pro-tumor activity of PTENα, while the F133/F263 residues of WDR5, essential for the interaction with PTENα/β and other WIN site ligands, are necessary but not the only requirement for the pro-tumor activity of WDR5. In fact, WDR5 also interacts with MYC, a well-known oncoprotein broadly overexpressed in some cancers, through its WBM site to promote tumorigenesis (Thomas et al. 2015). As PTENα and PTENβ share the same SSSRRSS fragment, our findings regarding PTENα are also applicable to PTENβ. Overall, our studies emphasize the role of the PTENα/β-WDR5 interaction in promoting oncogenic processes and provide a structural basis for development potential therapeutic targets for cancer treatment. Additionally, our studies suggest that the combined use of inhibitors targeting both the WIN site and the WBM site of WDR5 would be more effective in suppressing tumor growth than using the individual site inhibitor, which aligns with a recent research (Han et al. 2023).

In addition to their intracellular localization, the PTENα/β proteins can also be released into the extracellular space, where they are cleaved by the Furin proteinase. This cleavage generates a long C-terminal fragment, which has been shown to have significant tumor-suppressive properties (Zhang et al. 2022). Unfortunately, in liver cancer tissues, the expression of Furin is low and the cleavage of PTENα/β is reduced. Therefore, a potential therapeutic strategy for liver cancer could involve combining a PTENα/β-WDR5 protein-protein interface inhibitor with an activator of Furin. Further research is needed to test this hypothesis.

In conclusion, our study has revealed a novel binding motif within PTENα/β-NTE interacting with the WIN site of WDR5 to control downstream histone methylation and increase tumorigenic genes expression, cell proliferation, and tumor growth. Disrupting the PTENα/β-WDR5 interaction by mutations of the key interacting residues attenuates these effects, which indicates that novel therapies for cancer treatment could be potentially developed by targeting the interaction between PTENα/β and WDR5. Additionally, inhibiting both the WIN and WBM sites of WDR5 may further improve therapeutic efficacy. Overall, our research advances our understanding of the molecular mechanisms underlying cancer biology and offers promising possibilities for therapeutic interventions.

## Methods

### Plasmids, cell line and antibodies

The *E. coli* expression vectors, pET28GST-LIC, pET28a-MHL and pET32a-LIC were constructed in our laboratory. All the primers used in this study were provided in Supplementary Table 1.

Antibodies used in this study are as follows: mouse monoclonal anti-β-tubulin antibody (Sigma-Aldrich, Cat# T4026, 1:5000), mouse monoclonal anti-FLAG M2 antibody (Sigma-Aldrich, Cat# A8592, 1:5000), rabbit monoclonal anti-HA antibody (Sigma-Aldrich, Cat# H6908, 1:1000), rabbit monoclonal anti-NOTCH3 antibody (Cell Signaling Technology, Cat# 5276, 1:1000), rabbit monoclonal anti-H3K4me3 antibody (Cell Signaling Technology, Cat# 9751, 1:1000), rabbit monoclonal anti-PTEN antibody (Cell Signaling Technology, Cat# 9559, 1:1000), rabbit monoclonal anti-AKT antibody (Cell Signaling Technology, Cat# 4691, 1:1000), rabbit monoclonal anti-p-AKT (S473) antibody (Cell Signaling Technology, Cat# 4060, 1:1000).

### Cell lines and cell culture

Human embryonic kidney 293T (HEK293T) cells and human liver cancer SMMC-7721 cells were purchased from ATCC and Cell Bank of the Chinese Academy of Sciences in Shanghai, respectively. All the cell lines used in this study were cultured in DMEM supplemented with 10% fetal bovine serum, and underwent authentication using the short tandem repeat (STR) profile method and tested negative for mycoplasma contamination by PCR.

### Protein expression and purification

The DNA fragments of WDR5 (residues 22-334 or 1-334) and PTENα-NTE (residues 1-173) were subcloned into a pET28a-MHL vector to generate N-terminal 6×His-TEV-tagged fusion protein. Additionally, WDR5 (residues 22-334) was subcloned into a modified pET28GST-LIC vector to generate N-terminal GST-6×His-tagged fusion protein. PTENα (residues 1-576) was subcloned into a pET32a-LIC vector to generate N-terminal Trax-6×His-S-TEV-tagged fusion protein. All the plasmids were constructed using seamless assembly cloning (ABclonal Technology, RK21020) and confirmed by sequencing (Azenta Life Sciences).

The recombinant proteins were overexpressed in *E. coli* BL21 (DE3) Codon plus RIL (Stratagene, 230280) at 15°C for 24 h under induction with 0.25 mM IPTG (isopropyl-β-D-thiogalactoside) at an OD_600_ value of 0.8. They were then purified using affinity chromatography on Ni-nitrilotriacetate resin (GE Healthcare, 17526802) followed by TEV protease treatment to remove the tag for ITC assays and crystallization. The buffer conditions for Ni-affinity chromatography were as follows: lysis buffer: 20 mM Tris-HCl, pH 7.5, 250 mM NaCl, 5% glycerol, and 5 mM β-mercaptoethanol; wash buffer: 20 mM Tris-HCl, pH 7.5, 1 M NaCl, and 40 mM imidazole; elution buffer: 20 mM Tris-HCl, pH 7.5, 250 mM NaCl, and 250 mM imidazole.

WDR5 (residues 22-334 or 1-334) and PTENα (residues 1-576) proteins were further purified using a Superdex200 gel-filtration column (GE Healthcare, 28989335) with a buffer containing 20 mM Tris-HCl, pH 7.5, 150 mM NaCl, and 1 mM DTT. PTENα-NTE (residues 1-173) was further purified by ion exchange chromatography (HiTrap SP HP column, GE Healthcare, 17115201) using buffers with 20 mM Tris-HCl, pH 7.5, 1 mM DTT, adding either 50 mM NaCl (low salt buffer) or 1 M NaCl (high salt buffer), followed by dialysis using a buffer containing 20 mM Tris-HCl, pH 7.5, and 150 mM NaCl (ITC buffer). All the mutations involved in this study were constructed using Fast Mutagenesis System Kit (Transgene, FM111-02) according to manufacturer’s instruction and confirmed by DNA sequencing. Mutants were overexpressed and purified as the wild-type constructs described above. All the proteins were concentrated using Amicon Ultra-15 Centrifugal Filter Units (Millipore Corporation, UFC901024).

### Isothermal titration calorimetry (ITC)

For the ITC measurement, concentrated proteins were diluted into the ITC buffer, while lyophilized peptides were dissolved in the same buffer, and the pH was adjusted by adding 2 M NaOH dropwise. Peptide concentrations were estimated based on their mass. All measurements were conducted in duplicate at 25°C, utilizing an iTC-200 (MicroCal, Inc.) microcalorimeter.

In the cell chamber, a protein with a concentration of 50 µM was placed, and peptides or proteins with a concentration of 750 µM in the syringe were injected into the cell chamber for 20 successive injections with a spacing of 150 seconds. Control experiments were carried out under identical conditions to determine the heat signals that resulted from injecting peptides or proteins into the buffer. Data were fitted using the single-site binding model within the Origin software package (MicroCal, Inc.).

### Protein crystallization

For the crystals of WDR5^22-334^-PTENα-NTE^1-173^, purified WDR5^22-334^ was mixed with PTENα-NTE^1-173^ at a molar ratio of 1:1 and with trypsin at a mass ratio of 1:1000 (trypsin: protein mixture) (Dong et al. 2007; Zhang et al. 2018). Then, the mixture was crystallized using the sitting-drop vapor diffusion method at 18°C by adding 0.5 μL of the protein mixture (8 mg/mL) with 0.5 μL of the reservoir solution. The complex of WDR5^22-334^-PTENα-NTE^1-173^ crystallized in a buffer containing 0.2 M lithium sulfate monohydrate, 0.1 M HEPES, pH 7.5, and 25% w/v polyethylene glycol 3,350. As for the crystals of WDR5^1-334^, purified WDR5^1-334^ protein was mixed with PTENα-NTE^116-148^ peptide at a molar ratio of 1:3, and crystallized in a buffer containing 0.1 M magnesium formate dihydrate and 15% w/v polyethylene glycol 3,350. Before flash-freezing crystals in liquid nitrogen, crystals were soaked in a cryoprotectant consisting of 85% reservoir solution and 15% glycerol.

### Data collection and structure determination

The diffraction data of the crystals were collected at beamline BL18U1 of the Shanghai Synchrotron Radiation Facility (SSRF) at 100K. Diffraction images were processed using XDS (Kabsch 2010)/POINTLESS (Evans 2011)/AIMLESS (Evans and Murshudov 2013). PHASER (McCoy et al. 2007) software was used for molecular replacement searches. COOT (Emsley et al. 2010) was used for interactive model building. Following initial restrained model refinement with REFMAC (Murshudov et al. 2011), the model was automatically rebuilt with ARP/wARP (Langer et al. 2008). Subsequent model refinement was carried out using REFMAC and PHENIX (Adams et al. 2010). The crystal structure of WDR5 (PDB entry: 2H9M) was used as a coordinate for molecular replacement. REFMAC was applied for restrained model refinement (Murshudov et al. 2011). Crystal diffraction data and refinement statistics for the structures were presented in Table 1. All the structural figures were generated using PyMOL.

### GST pulldown assay

The purified GST-tagged fusion protein WDR5^22-334^ (100 μg) was bound to Glutathione Sepharose 4B (GE Healthcare, 28952360) for 1 h at 4°C. After washing with a buffer containing 20 mM Tris-HCl, pH 7.5, 150 mM NaCl, and 0.1% Triton X-100 for three times, the bound GST-tagged fusion proteins were incubated with purified tag-removed recombinant PTENα-NTE^1-173^ or full-length PTENα^1-576^, along with their respective mutants (300 μg). This incubation was carried out overnight at 4°C. Following another round of washing with the same buffer, the pulldown samples were eluted by adding 1×SDS protein loading buffer and loaded onto SDS-polyacrylamide gels and analyzed by coomassie brilliant blue (CBB) staining as previously described (Liu et al. 2017).

### Co-IP and western blotting

Eukaryotic expression plasmids, including 3×FLAG-WDR5, HA-PTENα, HA-PTENα-5A, and HA-PTENα-7A, were individually or co-transfected into HEK293T cells using Lipofectamine 2000 (ThermoFisher Scientific, 11668027) following the manufacturer’s protocol. The cells were harvested and lysed using IP (immunoprecipitation) lysis buffer containing 50 mM HEPES, pH 7.5, 150 mM NaCl, 10% glycerol, 1% Triton X-100, 1.5 mM MgCl_2_, and 1× protease inhibitor mixture (MCE, 180528) 48 h post-transfection as a previously described method with minor modifications (Wang et al. 2017; Yu et al. 2021). Briefly, the lysates were sonicated with five cycles of 0.3 s/0.7 s per mL and incubated for 30 min at 4°C and centrifuged at 12,000 rpm for 10 min at 4°C. The resulting supernatant were incubated with 20 μL of Anti-FLAG-Affinity-Gel (Selleck.cn, B23101) overnight at 4°C, followed by three washes with wash buffer (ITC buffer + 0.5% Triton X-100), each for 10 min. Then, 50 μL of 1× SDS protein loading buffer was added, and the mixture was boiled for 10 min. Subsequently, it was centrifuged at 12,000 rpm for 10 min, and the supernatant was removed for further analysis.

Input and IP protein samples were separated on a 4-12% Bis-Tris protein gel (GenScript, M41215C) using Tris-MOPS running buffer and then transferred onto a PVDF membrane and blocked overnight in 3% BSA in PBST buffer (PBS + 0.1% Tween 20). The membrane was then incubated with primary antibodies targeting the intended proteins for 1 h, followed by three 10-min washes in PBST. This process was repeated with secondary antibodies. Finally, the membrane was visualized using an Odyssey® CLx Imaging System (LI-COR).

### Quantitative real-time polymerase chain reaction (qRT-PCR)

Total RNA from various cell samples was extracted using TRIzol Reagent (Vazyme, R401-01). Subsequently, 1 μg of RNA was transcribed into complementary DNA (cDNA) using the ABScript III RT-PCR kit (ABclonal Technology, RK20429). The qPCR was performed using an SYBR Green reaction mix (ABclonal Technology, RK21203) with a LightCycler 96 System (Roche, Basel, Switzerland). Relative expression levels of target genes were calculated utilizing the 2^-ΔΔCt^ method (Ct, cycle threshold).

### Lentivirus-mediated WDR5/PTEN3 knockout

To generate *PTEN* or *WDR5* knockout in SMMC-7721 cell line, the following protocol was utilized: Cells were transfected with LentiCRISPR v2 plasmids containing specific sgRNAs: *PTEN* targeting sequence (sgRNA: ACAAAAGGAGATATCAAGAGG) or *WDR5* targeting sequence (sgRNA: TCTGAGTGGCGGATGACGAA). Subsequently, cells were subjected to puromycin selection. After selection, cells were diluted, and individual colonies were isolated. The sgRNA knockout efficacy was assessed by western blot analysis. PCR and DNA sequencing were used to confirm homozygous gene locus editing.

### Lentiviral transduction

Lentivirus was generated by co-transfecting HEK293T cells with the lentiviral construct pCMV-dR8.91 (Δ8.9) plasmid and the pMDG envelope-expressing plasmid using X-tremeGENE 9 DNA Transfection Reagent (Roche, 6365779001). Viral supernatant was collected within 24-48 h post-transfection for subsequent infection of the target cells.

### CCK-8 and colony formation assays

For CCK-8 assays, 2×10^3^ SMMC-7721 cells from different groups were seeded into each well of a 96-well plate. Cell cultures were established at 0 h (3 h after cells seeded was denoted as 0 h), 24 h, 48 h, 72 h, and 96 h. At each time point, 10 μL of CCK8 reagent (Selleck.cn, B34302) was added to each well. After 3 h of incubation, the optical density (OD) value was measured at 450 nm. Colony formation assays were performed by adding a total of 1000 SMMC-7721 cells to each group. After 10 days of culture, the colonies were fixed, stained, and counted by ImageJ.

### Mouse studies

A total of 1×10^6^ cells suspended in 100 μL of serum-free media were subcutaneously implanted into female nude mice aged 4 to 6 weeks. Tumor volumes were regularly monitored using calipers and calculated using the formula: length × (width)^2^/2. In accordance with animal care and ethical guidelines, the largest subcutaneous tumor mass on one flank was maintained below 1 cm^3^. All animal care and experimental procedures were conducted in strict compliance with ethical regulations governing animal research and were approved by the committee for the humane treatment of animals at Shanghai Jiao Tong University School of Medicine.

### Data availability

The coordinates and structure factors of this study were deposited in the Protein Data Bank (PDB) with accession codes 8X3S and 8X3R for the complex of WDR5-PTENα-NTE and WDR5 ligand-free structure, respectively. All other relevant data supporting the key findings of this study were available within the article and its supplementary information file or from the corresponding authors upon reasonable request.

## Acknowledgments

We would like to thank staffs of SSRF beamline BL18U1 for assistance in data collection. This work was supported by the National Natural Science Foundation of China grants (32271309), the Priority Academic Program Development of the Jiangsu Higher Education Institutes, China (PAPD), and Six talent Professorship of Jiangsu province, China (SWYY-104).

## Author contributions

X. H. purified and crystallized the protein; C. Z. conducted the animal experiments under the supervision of S. S.; X. S. determined the structure; Y. C. and Q. X. conducted the ITC assays with the help of Z. W.; X. H. cloned all the constructs and conducted the cellular study under the supervision of G. W., X. Z. and G. X.; J. M. reviewed the crystallographic model and provided all the vector plasmids; Y. L. conceived and designed the study, and wrote the paper with substantial contributions from all the other authors. All authors contributed to data analysis and approved the final version of the manuscript.

## References

Adams PD, Afonine PV, Bunkóczi G, Chen VB, Davis IW, Echols N, Headd JJ, Hung LW, Kapral GJ, Grosse-Kunstleve RW et al. 2010. PHENIX: a comprehensive Python-based system for macromolecular structure solution. Acta Crystallogr D Biol Crystallogr 66: 213–221.

Ali SA, Justilien V, Jamieson L, Murray NR, Fields AP. 2016. Protein kinase ciota drives a NOTCH3-dependent stem-like phenotype in mutant KRAS lung adenocarcinoma. Cancer Cell 29: 367–378.

Couture JF, Collazo E, Trievel RC. 2006. Molecular recognition of histone H3 by the WD40 protein WDR5. Nat Struct Mol Biol 13: 698–703.

Cui Z, Li H, Liang F, Mu C, Mu Y, Zhang X, Liu J. 2018. Effect of high WDR5 expression on the hepatocellular carcinoma prognosis. Oncol Lett 15: 7864–7870.

Dai X, Guo W, Zhan C, Liu X, Bai Z, Yang Y. 2015. WDR5 expression is prognostic of breast cancer outcome. PLoS One 10: e0124964.

Dharmarajan V, Lee JH, Patel A, Skalnik DG, Cosgrove MS. 2012. Structural basis for WDR5 interaction (Win) motif recognition in human SET1 family histone methyltransferases. J Biol Chem 287: 27275–27289.

Diao Y, Guo X, Li Y, Sun K, Lu L, Jiang L, Fu X, Zhu H, Sun H, Wang H et al. 2012. Pax3/7BP is a Pax7- and Pax3-binding protein that regulates the proliferation of muscle precursor cells by an epigenetic mechanism. Cell stem cell 11: 231–241.

Dias J, Van Nguyen N, Georgiev P, Gaub A, Brettschneider J, Cusack S, Kadlec J, Akhtar A. 2014. Structural analysis of the KANSL1/WDR5/KANSL2 complex reveals that WDR5 is required for efficient assembly and chromatin targeting of the NSL complex. Genes Dev 28: 929–942.

Dong A, Xu X, Edwards AM, Chang C, Chruszcz M, Cuff M, Cymborowski M, Di Leo R, Egorova O, Evdokimova E et al. 2007. In situ proteolysis for protein crystallization and structure determination. Nat Methods 4: 1019–1021.

Emsley P, Lohkamp B, Scott WG, Cowtan K. 2010. Features and development of Coot. Acta Crystallogr D Biol Crystallogr 66: 486–501.

Evans PR. 2011. An introduction to data reduction: space-group determination, scaling and intensity statistics. Acta Crystallogr D Biol Crystallogr 67: 282–292.

Evans PR, Murshudov GN. 2013. How good are my data and what is the resolution? Acta Crystallogr D Biol Crystallogr 69: 1204–1214.

Gagnon KT, Corey DR. 2012. Argonaute and the nuclear RNAs: new pathways for RNA-mediated control of gene expression. Nucleic Acid Ther 22: 3–16.

Ge Z, Song EJ, Kawasawa YI, Li J, Dovat S, Song C. 2016. WDR5 high expression and its effect on tumorigenesis in leukemia. Oncotarget 7: 37740–37754.

Guarnaccia AD, Rose KL, Wang J, Zhao B, Popay TM, Wang CE, Guerrazzi K, Hill S, Woodley CM, Hansen TJ et al. 2021. Impact of WIN site inhibitor on the WDR5 interactome. Cell Rep 34: 108636.

Guarnaccia AD, Tansey WP. 2018. Moonlighting with WDR5: a cellular multitasker. J Clin Med 7: 21.

Han QL, Zhang XL, Ren PX, Mei LH, Lin WH, Wang L, Cao Y, Li K, Bai F. 2023. Discovery, evaluation and mechanism study of WDR5-targeted small molecular inhibitors for neuroblastoma. Acta Pharmacol Sin 44: 877–887.

Hopkins BD, Fine B, Steinbach N, Dendy M, Rapp Z, Shaw J, Pappas K, Yu JS, Hodakoski C, Mense S et al. 2013. A secreted PTEN phosphatase that enters cells to alter signaling and survival. Science 341: 399–402.

Jeong JY, Kang H, Kim TH, Kim G, Heo JH, Kwon AY, Kim S, Jung SG, An HJ. 2017. MicroRNA-136 inhibits cancer stem cell activity and enhances the anti-tumor effect of paclitaxel against chemoresistant ovarian cancer cells by targeting Notch3. Cancer Lett 386: 168–178.

Jiang D, Gu X, He Y. 2009. Establishment of the winter-annual growth habit via FRIGIDA-mediated histone methylation at FLOWERING LOCUS C in Arabidopsis. Plant Cell 21: 1733–1746.

Ka M, Kim HG, Kim WY. 2022. WDR5-HOTTIP histone modifying complex regulates neural migration and dendrite polarity of pyramidal neurons via reelin signaling. Mol Neurobiol 59: 5104–5120.

Kabsch W. 2010. XDS. Acta Crystallogr D Biol Crystallogr 66: 125–132.

Kim JY, Banerjee T, Vinckevicius A, Luo Q, Parker JB, Baker MR, Radhakrishnan I, Wei JJ, Barish GD, Chakravarti D. 2014. A role for WDR5 in integrating threonine 11 phosphorylation to lysine 4 methylation on histone H3 during androgen signaling and in prostate cancer. Mol Cell 54: 613–625.

Kim JY, Yu J, Abdulkadir SA, Chakravarti D. 2016. KAT8 regulates androgen signaling in prostate cancer cells. Mol Endocrinol 30: 925–936.

Langer G, Cohen SX, Lamzin VS, Perrakis A. 2008. Automated macromolecular model building for X-ray crystallography using ARP/wARP version 7. Nat Protoc 3: 1171–1179.

Li DM, Sun H. 1997. TEP1, encoded by a candidate tumor suppressor locus, is a novel protein tyrosine phosphatase regulated by transforming growth factor beta. Cancer Res 57: 2124–2129.

Li J, Yen C, Liaw D, Podsypanina K, Bose S, Wang SI, Puc J, Miliaresis C, Rodgers L, McCombie R et al. 1997. PTEN, a putative protein tyrosine phosphatase gene mutated in human brain, breast, and prostate cancer. Science 275: 1943–1947.

Liang H, Chen X, Yin Q, Ruan D, Zhao X, Zhang C, McNutt MA, Yin Y. 2017. PTENβ is an alternatively translated isoform of PTEN that regulates rDNA transcription. Nat Commun 8: 14771.

Liang H, He SM, Yang JY, Jia XY, Wang P, Chen X, Zhang Z, Zou XJ, McNutt MA, Shen WH et al. 2014. PTEN α, a PTEN isoform translated through alternative initiation, regulates mitochondrial function and energy metabolism. Cell Metab 19: 836–848.

Liu Y, Qin S, Lei M, Tempel W, Zhang Y, Loppnau P, Li Y, Min J. 2017. Peptide recognition by heterochromatin protein 1 (HP1) chromoshadow domains revisited: Plasticity in the pseudosymmetric histone binding site of human HP1. J Biol Chem 292: 5655–5664.

Maehama T, Dixon JE. 1998. The tumor suppressor, PTEN/MMAC1, dephosphorylates the lipid second messenger, phosphatidylinositol 3,4,5-trisphosphate. J Biol Chem 273: 13375–13378.

Malaney P, Uversky VN, Dave V. 2013. The PTEN long N-tail is intrinsically disordered: increased viability for PTEN therapy. Mol Biosyst 9: 2877–2888.

McCoy AJ, Grosse-Kunstleve RW, Adams PD, Winn MD, Storoni LC, Read RJ. 2007. Phaser crystallographic software. J Appl Crystallogr 40: 658–674.

Mo R, Rao SM, Zhu YJ. 2006. Identification of the MLL2 complex as a coactivator for estrogen receptor alpha. J Biol Chem 281: 15714–15720.

Murshudov GN, Skubák P, Lebedev AA, Pannu NS, Steiner RA, Nicholls RA, Winn MD, Long F, Vagin AA. 2011. REFMAC5 for the refinement of macromolecular crystal structures. Acta Crystallogr D Biol Crystallogr 67: 355–367.

Myers MP, Stolarov JP, Eng C, Li J, Wang SI, Wigler MH, Parsons R, Tonks NK. 1997. P-TEN, the tumor suppressor from human chromosome 10q23, is a dual-specificity phosphatase. Proc Natl Acad Sci U S A 94: 9052–9057.

Patel A, Vought VE, Dharmarajan V, Cosgrove MS. 2008. A conserved arginine-containing motif crucial for the assembly and enzymatic activity of the mixed lineage leukemia protein-1 core complex. J Biol Chem 283: 32162–32175.

Qin S, Liu Y, Tempel W, Eram MS, Bian C, Liu K, Senisterra G, Crombet L, Vedadi M, Min J. 2014. Structural basis for histone mimicry and hijacking of host proteins by influenza virus protein NS1. Nat Commun 5: 3952.

Ravnskjaer K, Hogan MF, Lackey D, Tora L, Dent SY, Olefsky J, Montminy M. 2013. Glucagon regulates gluconeogenesis through KAT2B and WDR5-mediated epigenetic effects. J Clin Invest 123: 4318–4328.

Ruthenburg AJ, Wang W, Graybosch DM, Li H, Allis CD, Patel DJ, Verdine GL. 2006. Histone H3 recognition and presentation by the WDR5 module of the MLL1 complex. Nat Struct Mol Biol 13: 704–712.

Sansal I, Sellers WR. 2004. The biology and clinical relevance of the PTEN tumor suppressor pathway. J Clin Oncol 22: 2954–2963.

Schuettengruber B, Martinez AM, Iovino N, Cavalli G. 2011. Trithorax group proteins: switching genes on and keeping them active. Nat Rev Mol Cell Biol 12: 799–814.

Schuetz A, Allali-Hassani A, Martin F, Loppnau P, Vedadi M, Bochkarev A, Plotnikov AN, Arrowsmith CH, Min J. 2006. Structural basis for molecular recognition and presentation of histone H3 by WDR5. EMBO J 25: 4245–4252.

Shen SM, Zhang C, Ge MK, Dong SS, Xia L, He P, Zhang N, Ji Y, Yang S, Yu Y et al. 2020. PTENα and PTENβ promote carcinogenesis through WDR5 and H3K4 trimethylation. Nat Cell Biol 22: 135–135.

Shilatifard A. 2012. The COMPASS family of histone H3K4 methylases: mechanisms of regulation in development and disease pathogenesis. Annu Rev Biochem 81: 65–95.

Song JJ, Kingston RE. 2008. WDR5 interacts with mixed lineage leukemia (MLL) protein via the histone H3-binding pocket. J Biol Chem 283: 35258–35264.

Steck PA, Pershouse MA, Jasser SA, Yung WK, Lin H, Ligon AH, Langford LA, Baumgard ML, Hattier T, Davis T et al. 1997. Identification of a candidate tumour suppressor gene, MMAC1, at chromosome 10q23.3 that is mutated in multiple advanced cancers. Nat Genet 15: 356–362.

Sulis ML, Parsons R. 2003. PTEN: from pathology to biology. Trends Cell Biol 13: 478–483.

Tan M, Li S, Juillard F, Chitas R, Custodio TF, Xue H, Szymula A, Sun Q, Liu B, Alvarez AL et al. 2021. MLL1 is regulated by KSHV LANA and is important for virus latency. Nucleic Acids Res 49: 12895–12911.

Tennant BR, Hurley P, Dhillon J, Gill A, Whiting C, Hoffman BG. 2015. The TrxG complex mediates cytokine induced de novo enhancer formation in islets. PLoS One 10: e0141470.

Thomas LR, Wang Q, Grieb BC, Phan J, Foshage AM, Sun Q, Olejniczak ET, Clark T, Dey S, Lorey S et al. 2015. Interaction with WDR5 promotes target gene recognition and tumorigenesis by MYC. Mol Cell 58: 440–452.

Wang JR, Sun PH, Ren ZX, Meltzer HY, Zhen XC. 2017. GSK-3β interacts with dopamine D1 receptor to regulate receptor function: implication for prefrontal cortical D1 receptor dysfunction in schizophrenia. CNS Neurosci Ther 23: 174–187.

Wang YY, Liu LJ, Zhong B, Liu TT, Li Y, Yang Y, Ran Y, Li S, Tien P, Shu HB. 2010. WDR5 is essential for assembly of the VISA-associated signaling complex and virus-triggered IRF3 and NF-kappaB activation. Proc Natl Acad Sci U S A 107: 815–820.

Wysocka J, Swigut T, Milne TA, Dou Y, Zhang X, Burlingame AL, Roeder RG, Brivanlou AH, Allis CD. 2005. WDR5 associates with histone H3 methylated at K4 and is essential for H3 K4 methylation and vertebrate development. Cell 121: 859–872.

Xu Z, Gao X, He Y, Ju J, Zhang M, Liu R, Wu Y, Ma C, Ma C, Lin Z et al. 2012. Synergistic effect of SRY and its direct target, WDR5, on Sox9 expression. PLoS One 7: e34327.

Yin Y, Shen WH. 2008. PTEN: a new guardian of the genome. Oncogene 27: 5443–5453.

Yu W, Wang B, Zhou L, Xu G. 2021. Endoplasmic reticulum stress-mediated p62 downregulation inhibits apoptosis via c-Jun upregulation. Biomol Ther (Seoul) 29: 195–204.

Zhang C, Ma HM, Dong SS, Zhang N, He P, Ge MK, Xia L, Yu JX, Xia Q, Chen GQ et al. 2022. Furin extracellularly cleaves secreted PTENα/β to generate C-terminal fragment with a tumor-suppressive role. Cell Death Dis 13: 532.

Zhang P, Lee H, Brunzelle JS, Couture JF. 2012. The plasticity of WDR5 peptide-binding cleft enables the binding of the SET1 family of histone methyltransferases. Nucleic Acids Res 40: 4237–4246.

Zhang Q, Liang H, Zhao X, Zheng L, Li Y, Gong J, Zhu Y, Jin Y, Yin Y. 2021. PTENε suppresses tumor metastasis through regulation of filopodia formation. EMBO J 40: e105806.

Zhang Y, Lei M, Yang X, Feng Y, Yang Y, Loppnau P, Li Y, Yang Y, Min J, Liu Y. 2018. Structural and histone binding studies of the chromo barrel domain of TIP60. FEBS Lett 592: 1221–1232.

Zhu ED, Demay MB, Gori F. 2008. WDR5 is essential for osteoblast differentiation. J Biol Chem 283: 7361–7367.

